# Targeted plasma proteomics uncover novel proteins associated with *KIF5A*-linked SPG10 and ALS spectrum disorders

**DOI:** 10.1101/2025.04.29.651203

**Authors:** Jarosław Dulski, Arun K. Boddapati, Barbara Risi, Pablo Iruzubieta, Antonio Orlacchio, Roberto Fernández-Torrón, Tamara Castillo-Triviño, Adolfo López de Munain, Steve Vucic, Laura Donker Kaat, Tahsin Stefan Barakat, Leonard Petrucelli, Mercedes Prudencio, John E. Landers, Jochen H. Weishaupt, Andreas Prokop, Massimiliano Filosto, Zbigniew K. Wszolek, Devesh C. Pant

## Abstract

KIF5A (Kinesin family member 5A) is a motor protein that functions as a key component of the axonal transport machinery. Variants in *KIF5A* are linked to several neurodegenerative diseases, mainly spastic paraplegia type 10 (SPG10), Charcot-Marie-Tooth disease type 2 (CMT2), and amyotrophic lateral sclerosis (ALS). These diseases share motor neuron involvement but vary significantly in clinical presentation, severity, and progression. *KIF5A* variants are mainly categorized into N-terminal variants associated with SPG10/CMT2 and C-terminal variants linked to ALS. This study utilized a novel multiplex NULISA targeted platform to analyze plasma proteome from *KIF5A*-linked SPG10, ALS patients and compared to healthy controls. Our results revealed distinct proteomic signatures, with significant alterations in proteins related to synaptic function, and inflammation. Notably, neurofilament light polypeptide, a biomarker for neurodegenerative diseases, was elevated in *KIF5A* ALS but not in SPG10 patients. Moreover, these findings can now be taken forward to gain mechanistic understanding of axonopathies linking to N-*vs* C-terminal *KIF5A* variants affecting both central and peripheral nervous systems.

## Introduction

The *KIF5A* gene encodes a motor protein that is part of the kinesin-1 family, responsible for facilitating the transport of cellular cargoes along microtubules.^1^ Several studies have reported variants in *KIF5A* (OMIM# 602821) as significant contributors to a range of neurodegenerative diseases, particularly spastic paraplegia type 10 (SPG10), Charcot-Marie-Tooth disease type 2 (CMT2), neonatal intractable myoclonus (NEIMY), and recently amyotrophic lateral sclerosis (ALS).^2-7^ KIF5A is the only motor protein within the KIF5 family (which includes KIF5B and KIF5C) that has been identified as mutated in neurodegenerative disorders with high penetrance. Although *KIF5A* linked neurodegenerative disorders share some pathophysiological and clinical features, primarily affecting motor neurons and leading to progressive impairment in motor function, they differ substantially in the distribution of symptoms, severity of symptoms and overall prognosis.^2, 3, 5, 6^

The clinical spectrum of *KIF5A* variants has expanded since the first identification of *KIF5A*-linked to SPG10. A large family pedigree initially identified an autosomal dominant locus for SPG10, which was subsequently mapped to chromosome 12 by Rubinsztein lab (UK).^6, 8^ SPG10 typically presents with progressive spasticity and weakness confined to the lower limbs, resulting from degeneration of the corticospinal tracts. Interestingly, a high degree of intra-familial phenotypic variability was initially reported in a *KIF5A* family where the index patient had SPG10 and father had axonal CMT with moderate lower limb amyotrophy, distal hypoesthesia and axonal neuropathy but no spasticity.^9^ The intra-familial phenotypic heterogeneity was further observed in an Asian *KIF5A* SPG10 family which was characterized by late-onset, dysarthria and fasciculations. The presence of fasciculations was never reported in SPG10 and the mutation is located at the N-terminal motor domain of KIF5A protein, where the previously reported SPG10, or rare CMT2 variants are clustered.^10^ Due to observed fasciculations, the authors suggested that *KIF5A* may need to be considered in the differential diagnosis of sporadic ALS. Other *KIF5A* rare phenotypes include adult-onset distal spinal muscular atrophy (SMA), reported in one family with variant in motor domain.^11^

*KIF5A* was previously identified as a potential ALS candidate gene, but the study lacked sufficient cases to make a definitive conclusion.^12^ In 2018, the group of Landers (USA) and Weishaupt (Germany) reported *KIF5A*-linked familial ALS variants in the C-terminal cargo domain of KIF5A protein.^2, 5^ The *KIF5A*-linked ALS patients display younger age at onset and longer survival. ALS typically involve both upper and lower motor neurons present with rapid-onset muscle weakness and wasting, while *KIF5A* related ALS tend to show more insidious progression. Additionally, ALS and frontotemporal dementia (FTD) has been reported in one large family with *KIF5A* cargo domain variant by Traynor lab (USA).^13^ Conversely, severe developmental syndrome NEIMY has been linked to *de novo* variants in the cargo domain of *KIF5A*. These patients typically presented with myoclonic seizures, accompanied by symptoms such as hypotonia, optic nerve abnormalities, dysphagia, apnea, early developmental stagnation, and progressive leukoencephalopathy.^4, 7^

Axonopathies caused by *KIF5A* variants have been classically associated with a slow progression of muscle weakness leading to severe disability. Many cell biologists have examined *KIF5A* mediated axonal transport by expressing the mutant motor proteins and binding partners using cellular models.^14-16^ These studies are informative but have certain limitations, as they may not fully capture the complexities of axonopathies in its natural context. Gaining a clearer understanding of how *KIF5A* variants contribute to the diverse clinical outcomes seen in various disorders, including axonopathy, is crucial for advancing our understanding of disease, improving patient counseling, and guiding future therapeutic approaches. Data from *in vitro* over-expression studies suggest that the majority of *KIF5A* motor domain variants linked to SPG10/CMT2, display reduce microtubule binding and KIF5A motility, while cargo domain variants linked to ALS, lead to hyperactive kinesin.^17-20^ Despite knowing those structure-functional correlations, it remains unclear how and why they link to such divergent phenotypes. Potential new understanding might arise from patient proteomics studies comparing potential changes that correlate with the different *KIF5A* variants.

In this study, we collected genetic, phenotypic and blood-plasma from an international cohort of patients with different variants in *KIF5A*, leveraging on a novel targeted proteomics platform using *KIF5A* patients’ plasma. Through this approach, we identified a set of novel targets that can now be taken forward to gain mechanistic understanding of axonopathies linking to N-*vs* C-terminal *KIF5A* variants.

## Methods

### Study participants

We reached out to different neuromuscular centers worldwide that had previously reported or following up with *KIF5A* patients. Plasma samples were collected from total n=19 *KIF5A* patients and n=18 aged-matched controls (**Table 1**). Out of nineteen patients, n=7 diagnosed with ALS, n=7 with SPG10, n=2 overlap with SPG10 (CMT2) and n=3 presymptomatic according to the El Escorial criteria.^21^ To maximize the potential of a broader *KIF5A* cohort, we defined two primary groups: one for N-terminal variants (associated with SPG10/CMT2) and another for C-terminal variants (linked to ALS). All participants provided informed consent and the ethics board granted approval (IRB#: 16-009414).

**Table 1.** Characterization of *KIF5A* cohort assessed in the study.

| Sample ID | Genetic variant | Protein change | Disease | Gender | Age |
| --- | --- | --- | --- | --- | --- |
| B_08 | c.2341A>G | p.Lys781Glu | SPG10/CMT2 | M | 54 |
| B_10 | c.839G>A | p.Arg280His | SPG10 | M | 37 |
| B_11 | c.1051_1053delGAG | p.Glu351del | CMT2 | F | 30 |
| C_10 | c.967C>T | p.Arg323Trp | SPG10 | F | 25 |
| C_11 | c.967C>T | p.Arg323Trp | SPG10 | F | 25 |
| D_01 | c.967C>T | p.Arg323Trp | SPG10 | F | 54 |
| D_02 | c.1329dupG | p.Lys444Glufs*38 | SPG10 | M | 26 |
| D_03 | c.611G>A | p.Arg204Gln | SPG10 | M | 63 |
| D_04 | c.1785delA | p.Lys595Asn*27 | SPG10 | M | 34 |
| C_05 | c.3020G>A | 3' splice junction | presymptomatic | M | 34 |
| C_06 | c.3020G>A | 3' splice junction | presymptomatic | M | 34 |
| C_09 | c.3020G>A | 3' splice junction | presymptomatic | F | 26 |
| B_09 | c.2764G>A | p.Val922Ile | ALS | M | 67 |
| C_01 | c.2993-1G>A | 5' splice junction | ALS | M | 45 |
| C_02 | c.3020G>A | 3' splice junction | ALS | M | 54 |
| C_03 | c.3020G>A | 3' splice junction | ALS | M | 54 |
| C_04 | c.3020G>A | 3' splice junction | ALS | M | 54 |
| C_07 | c.3020G>A | 3' splice junction | ALS | F | 49 |
| C_08 | c.3020G>A | 3' splice junction | ALS | M | 59 |

M, male; F, female; SPG10, spastic paraplegia-10; CMT2, Charcot-Marie-Tooth disease type 2; ALS, amyotrophic lateral sclerosis

### Blood collection

Standard venipuncture was conducted to collect 10 ml venous blood in an EDTA tube for plasma separation. Blood was centrifuged at 2,465*g* for 15 min at 4°C. Samples were processed within 30 min of collection. Plasma was aliquoted into 2 ml screw-cap tubes and stored at −80°C.

### Proteomics platform

The Nucleic Acid Linked Immuno-Sandwich Assay (NULISA) was carried out at Alamar Biosciences (Fremont, CA) as described previously.^22^ Briefly, plasma samples were initially stored at −80°C. After thawing on ice, the samples were re-centrifuged at 10,000*g* for 10 min. A 10 μL portion of the supernatant was transferred to a 96-well plate and analyzed using Alamar’s NULISASeq™ CNS Disease Panel (**Table 2**). NULISA’s design features enhance both multiplexing and automation. It creates the reporter DNA molecule via ligation, which simplifies the design of assays with multiple targets. The NULISASeq process was automated, beginning with the formation of immunocomplexes through DNA-barcoded capture and detection antibodies. These immunocomplexes were captured on paramagnetic oligo-dT beads, washed, and then released into a low-salt buffer, followed by capture, and washing on streptavidin beads. The DNA strands at the ends of the immunocomplexes were ligated to form a reporter molecule containing both target-specific and sample-specific barcodes. The capture antibody is attached to a partially double-stranded DNA, which contains a poly-A tail and a target-specific molecular identifier (TMI), while the detection antibody is linked to another partially double-stranded DNA, featuring a biotin group and a complementary TMI. These reporter molecules were then pooled, amplified via PCR, purified, and sequenced using the Illumina NextSeq 2000.

**Table 2.** List of NULISA protein panel used in the study.

| **Target** | **Uniprot ID** | **Protein name** |
| --- | --- | --- |
| ACHE | P22303 | Acetylcholinesterase |
| AGRN | O00468 | Agrin |
| ANXA5 | P08758 | Annexin A5 |
| APOE | P02649 | Apolipoprotein E |
| ARSA | P15289 | Arylsulfatase A |
| Aβ38 | P05067 | Amyloid-beta precursor protein |
| Aβ40 | P05067 | Amyloid-beta precursor protein |
| Aβ42 | P05067 | Amyloid-beta precursor protein |
| BACE1 | P56817 | Beta-secretase 1 |
| BASP1 | P80723 | Brain abundant membrane attached signal protein 1 |
| BDNF | P23560 | Brain-derived neurotrophic factor |
| CCL11 | P51671 | Eotaxin |
| CCL13 | Q99616 | C-C motif chemokine 13 |
| CCL17 | Q92583 | C-C motif chemokine 17 |
| CCL2 | P13500 | C-C motif chemokine 2 |
| CCL22 | O00626 | C-C motif chemokine 22 |
| CCL26 | Q9Y258 | C-C motif chemokine 26 |
| CCL3 | P10147 | C-C motif chemokine 3 |
| CCL4 | P13236 | C-C motif chemokine ligand 4 |
| CD40LG | P29965 | CD40 ligand |
| CD63 | P08962 | CD63 antigen |
| CHI3L1 | P36222 | Chitinase-3-like protein 1 |
| CHIT1 | Q13231 | Chitotriosidase-1 |
| CNTN2 | Q02246 | Contactin-2 |
| CRH | P06850 | Corticoliberin |
| CRP | P02741 | C-reactive protein |
| CSF2 | P04141 | Granulocyte-macrophage colony-stimulating factor |
| CST3 | P01034 | Cystatin-C |
| CX3CL1 | P78423 | Fractalkine |
| CXCL1 | P09341 | Growth-regulated alpha protein |
| CXCL10 | P02778 | C-X-C motif chemokine 10 |
| CXCL8 | P10145 | Interleukin-8, IL8 |
| ENO2 | P09104 | Gamma-enolase |
| FABP3 | P05413 | Fatty acid-binding protein, heart |
| FCN2 | Q15485 | Ficolin-2 |
| FGF2 | P09038 | Fibroblast growth factor 2 |
| FLT1 | P17948 | Vascular endothelial growth factor receptor 1 |
| FOLR1 | P15328 | Folate receptor alpha |
| GDF15 | Q99988 | Growth/differentiation factor 15 |
| GDNF | P39905 | Glial cell line-derived neurotrophic factor |
| GFAP | P14136 | Glial fibrillary acidic protein |
| GOT1 | P17174 | Aspartate aminotransferase, cytoplasmic |
| HBA1 | P69905 | Hemoglobin subunit alpha |
| HTT | P42858 | Huntingtin |
| ICAM1 | P05362 | Intercellular adhesion molecule 1 |
| IFNG | P01579 | Interferon gamma |
| IGF1R | P08069 | Insulin-like growth factor 1 receptor |
| IGFBP7 | Q16270 | Insulin-like growth factor-binding protein 7 |
| IL10 | P22301 | Interleukin-10 |
| IL12p70 | P29459\|P29460 | Interleukin-12 subunit beta\|Interleukin-12 subunit alpha |
| IL13 | P35225 | Interleukin-13 |
| IL15 | P40933 | Interleukin-15 |
| IL16 | Q14005 | Pro-interleukin-16 |
| IL18 | Q14116 | Interleukin-18 |
| IL2 | P60568 | Interleukin-2 |
| IL33 | O95760 | Interleukin 33 |
| IL5 | P05113 | Interleukin-5 |
| IL6 | P05231 | Interleukin-6 |
| IL6R | P08887 | Interleukin-6 receptor subunit alpha |
| IL7 | P13232 | Interleukin-7 |
| IL9 | P15248 | Interleukin-9 |
| KDR | P35968 | Vascular endothelial growth factor receptor 2 |
| KLK6 | Q92876 | Kallikrein-6 |
| MAPT | P10636 | Microtubule-associated protein tau |
| MDH1 | P40925 | Malate dehydrogenase, cytoplasmic |
| MME | P08473 | Membrane metalloendopeptidase |
| MSLN | Q13421 | Mesothelin |
| NEFH | P12036 | Neurofilament heavy polypeptide |
| NEFL | P07196 | Neurofilament light polypeptide |
| NGF | P01138 | Beta-nerve growth factor |
| NPTX1 | Q15818 | Neuronal pentraxin-1 |
| NPTX2 | P47972 | Neuronal pentraxin-2 |
| NPTXR | O95502 | Neuronal pentraxin receptor |
| NPY | P01303 | Neuropeptide Y |
| NRGN | Q92686 | Neurogranin |
| PDGFRB | P09619 | Platelet-derived growth factor receptor beta |
| PGF | P49763 | Placenta growth factor |
| PGK1 | P00558 | Phosphoglycerate kinase 1 |
| POSTN | Q15063 | Periostin |
| PRDX6 | P30041 | Peroxiredoxin-6 |
| PSEN1 | P49768 | Presenilin 1 |
| pSNCA-129 | P37840 | Alpha-synuclein |
| pTau-181 | P10636 | Microtubule-associated protein tau |
| pTau-231 | P10636 | Microtubule-associated protein tau |
| REST | Q13127 | RE1 silencing transcription factor |
| RUVBL2 | Q9Y230 | RuvB like AAA ATPase 2 |
| S100A12 | P80511 | Protein S100-A12 |
| S100B | P04271 | S100 calcium binding protein B |
| SAA1 | P0DJI8 | Serum amyloid A-1 protein |
| SFRP1 | Q8N474 | Secreted frizzled-related protein 1 |
| SFTPD | P35247 | Pulmonary surfactant-associated protein D |
| SLIT2 | O94813 | Slit homolog 2 protein |
| SMOC1 | Q9H4F8 | SPARC-related modular calcium-binding protein 1 |
| SNAP25 | P60880 | Synaptosomal-associated protein 25 |
| SNCA | P37840 | Alpha-synuclein |
| SOD1 | P00441 | Superoxide dismutase [Cu-Zn] |
| SQSTM1 | Q13501 | Sequestosome-1 |
| TAFA5 | Q7Z5A7 | Chemokine-like protein TAFA-5 |
| TARDBP | Q13148 | TAR DNA-binding protein 43 |
| TEK | Q02763 | Angiopoietin-1 receptor |
| TIMP3 | P35625 | Metalloproteinase inhibitor 3 |
| TNF | P01375 | Tumor necrosis factor |
| TREM1 | Q9NP99 | Triggering receptor expressed on myeloid cells 1 |
| TREM2 | Q9NZC2 | Triggering receptor expressed on myeloid cells 2 |
| VCAM1 | P19320 | Vascular cell adhesion protein 1 |
| VEGFA | P15692 | Vascular endothelial growth factor A |
| VEGFD | O43915 | Vascular endothelial growth factor D |
| VGF | O15240 | VGF nerve growth factor inducible |
| VSNL1 | P62760 | Visinin-like protein 1 |

### Data normalization and quality control

The sequencing data were processed using the NULISAseq algorithm developed by Alamar Biosciences (CA).^22^ In brief, barcodes corresponding to both the sample and target were quantified. For normalization, the count for each target was divided by the internal control counts for that well. The resulting data were rescaled, incremented by 1, and log_2_ transformed to derive the NULISA Protein Quantification (NPQ) arbitrary unit, which was then used for subsequent statistical analysis. Moreover, inter-plate normalization is part of the standard normalization process and is carried out to make data comparable across runs.

### Statistical Analysis

Data analyses were performed using R software (version 4.4.1, the R foundation for statistical analysis). NPQ values from NULISA were already log_2_ transformed. Normalized NPQ values have been used to perform differential expression (DE) analysis by fitting linear model (lm, a default package of R package) for each protein using a reduced model representing only two comparisons. The DE is a statistical approach to detect and identify biological targets whose expression varies between groups. We eliminated proteins whose NPQ values were below the NULISA limit of detection, *p*-value < 0.05 was considered statistically significant. Our study focuses on a predefined set of proteins, *p*-values are presented as independent results without adjustments. Additionally, we check for individual protein performance evaluation via bootstrapped AUC calculation on the *p* < 0.05 significant proteins across all the different comparisons (**Table S1, Figure S2**). For each protein and comparison group, we performed 1000 bootstrap iterations. In each iteration, we sampled with replacement from the original dataset, maintaining the same sample size.

## Results

The main goal of our study was to identify proteins that exhibit alterations in individuals carrying autosomal dominant *KIF5A* variants, primarily associated with SPG10 or ALS, with a view to improve our understanding of the mechanisms that determine such different disease outcomes. Blood is frequently utilized in liquid biopsy for diagnostic purposes. However, the blood proteome, includes proteins actively secreted into the bloodstream, as well as the proteomes from other tissues holds even greater potential in providing a real-time insight into disease. To do this, we used multiplexed NULISA with next-generation sequencing readouts for detecting central nervous system targets. We analyzed 109 unique proteins which passed the NULISA limit of detection, from human *KIF5A* mutation carriers and control plasma samples (**Figure 1**). We identified 109 proteins in the plasma of control and disease groups to pinpoint potential significant differences. Participants in the control group were sex and age matched to the patient group, thus further enhancing the power of our study to detect reliable proteomic alterations.

**Figure 1.**
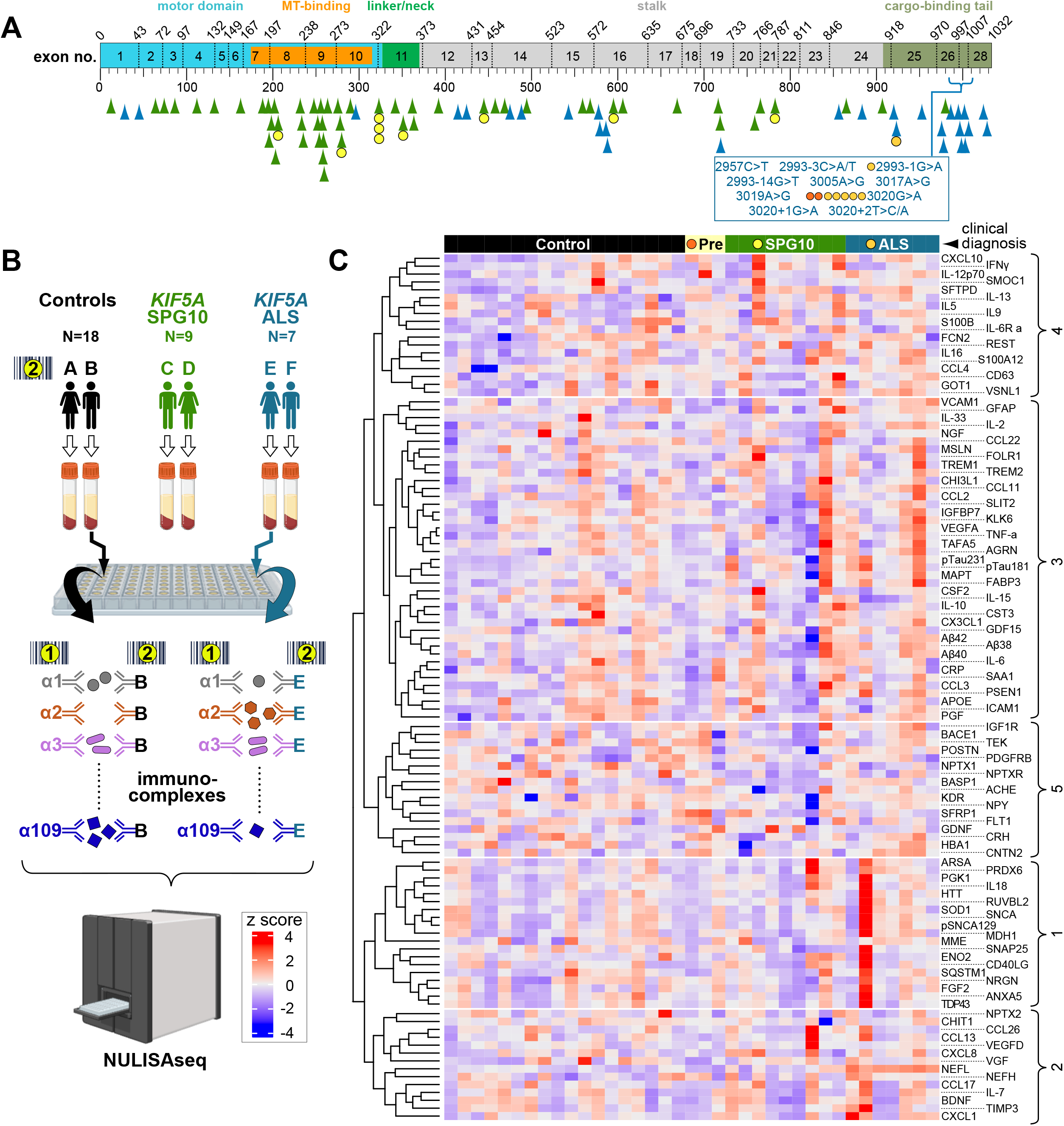
Design and outcome of the *KIF5A* linked SPG10, ALS plasma proteomics study. (**A**) Schematic representation of the *KIF5A* isoform 202 (ENST00000455537.7, 5779bp, 1032aa), with functional domains colour-coded as indicated, angularly numbered stippled vertical lines showing residue position of exon borders (number of every other exon provided); point mutations with identified residue changes are indicated by arrow heads, data taken from^2, 5, 34, 35, 58-60^ (and this study; see Table 1): green arrow heads represent the SPG10/CMT2-linked mutations: V12A, Y63C, D73N, Q87E, R111P, K132R R162W/P, S189P, R191H, A194P, T196N, M198T, S202N, S203C, <u>R204Q/W/P</u>, V231L D232N/W, G235E, L249V, E251K, K253N, L255M, N256Δ/S, K257N, S258L, L259Q, L262P A268T, Y276C, P278L, <u>R280H</u>/C/L, D290H, <u>R323W</u>, Q341R, <u>K444Efs*38</u>, <u>E351Δ</u>, A361V, K362N, A391V, S458F, R468W, L494M, L558P, G568R, <u>K595N*27</u>, R606W, A669T, R716W, R718W, E758K, Q764*, <u>K781E</u>, R864*, K907M, A980V; blue arrow heads represent the ALS-linked mutations: K29R, P46S, R297Q, E413G, R423H, Q474H, L488R, H542N, S577G, A579T, T585A, R588Q, R716Q, D853N, E881K, K920fs, <u>V922I</u>, T976E, Q952fs, C975fs, S978L/fs, P986L, D996fs, N997fs, N999fs/Δ, T1001Qfs, D1002G, R1007G/K, F1023C, E1028D; C-terminal ALS-linked intronic and exonic point mutations occurring only at DNA level are listed in the blue box; underlined mutations indicate patients of this study, also represented as circles where each circle indicates an individual patient. (**B**) Experimental flow of the blood analysis: antibodies detecting 109 biomarker proteins (α1-α109) are DNA-barcoded either to indicate the antibody identity (‘1’ in yellow circle) or the patient’s identity (‘2’ in yellow circle); readily formed immuno-complexes with both sets of antibodies from each blood sample are combined and subjected to NULISAseq analysis quantifying the abundance of each barcode pair. (**C**) Heatmap displaying the log_2_-normalized read counts (z score: from -4, blue, lower expression to 4, red, higher expression). Rows represent individual proteins indicated on the right (for abbreviations see Tab.2) clustered based on similarity, while columns represent individual patient samples clustered by clinical diagnosis (color-coded on top). The dendrogram on the left indicates hierarchical clustering of proteins (5 distinct clusters; indicated by curly brackets on the right), revealing distinct patterns of expression across clinical diagnoses.

Plasma analyses of *KIF5A*-linked ALS patients showed a significant increase in a range of proteins that could be grouped under the term: amyloidopathy (amyloid-beta 40 [Aβ38]), microtubule stabilization (pTau-181, or pTau-231), synaptic function (contactin-2 [CNTN2], presenilin-1 [PSEN1]), fatty acid metabolism regulation (fatty acid binding protein 3 [FABP3]), cell cycle regulation (RuvB-like 2 [RUVBL2]), inflammation regulation (C X-C Motif Chemokine Ligand 1 [CXCL1], and pulmonary surfactant-associated protein D [SFTPD]). Neuronal pentraxin-1 (NPTX1), interleukin-15 [IL15] were the only two proteins that displayed significant downregulation in *KIF5A* ALS (**Figure 2**). The most notable finding in the plasma of *KIF5A*-linked ALS patients was neurofilament light polypeptide (NEFL), a well-known biomarker for neurodegenerative diseases that has been extensively studied in the context of ALS.^23-25^ Further analysis conducted on three presymptomatic individuals carrying a *KIF5A* splice site mutation (c.3020G>A) known to have strong links to ALS ^5, 26^ revealed significant deviations from *KIF5A* ALS patients: the NEFL and Aβ40 proteins that were found significantly increased in the main analyses were found to be significantly reduced in presymptomatic carriers (**Figure S1**).

**Figure 2.**
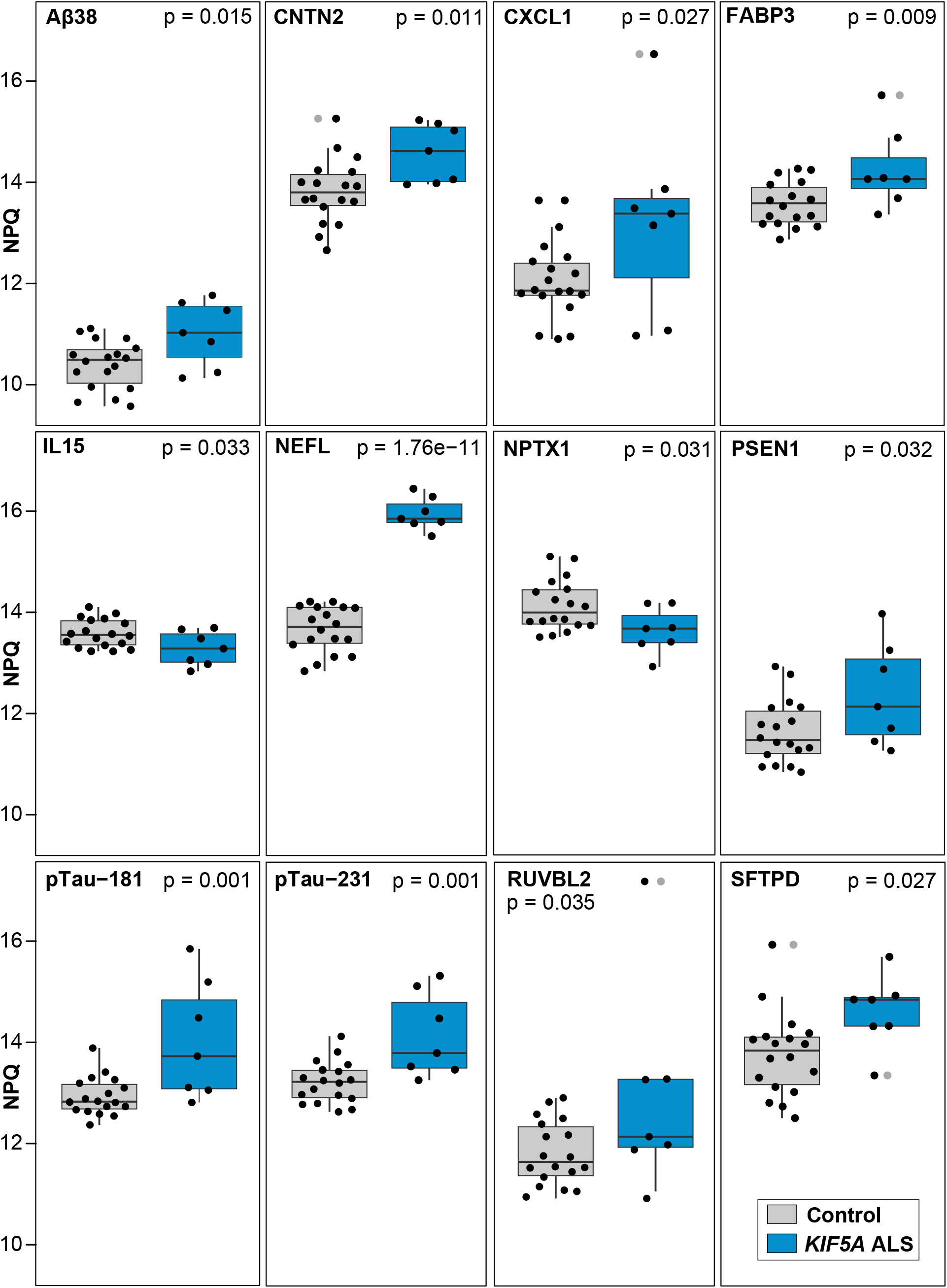
Comparison of NULISAseq biomarkers between control and *KIF5A*-linked ALS. Box plots illustrating the distributions of targets in plasma of controls (n=18) and *KIF5A* ALS (n=7) participants with significant or marginally significant differences for proteins (Aβ38, CNTN2, FABP3, NEFL, NPTX1, pTau-181, pTau-231, and SFTPD). The y-axis represents log_2_-tranformation of normalized read counts (NULISA Protein Quantification, NPQ). On each box, the central mark indicates the median, and the bottom and top edges of the box indicate the 25th and 75th percentiles, respectively. The whiskers extend to the most extreme data points not considered outliers, and the outliers are plotted individually using the grey dot symbol. The *p* values were determined using linear model. Significance determination was based on *p*-value < 0.05.

Plasma analyses of *KIF5A*-linked SPG10 patients revealed significant decrease in three unique proteins which include the actin-cytoskeleton protein BASP1 (brain acid soluble protein 1), the synaptic function and plasticity regulator NPTXR (neuronal pentraxin receptor), the synaptic protein SNAP25 (synaptosomal-associated protein 25), whereas the common vascular inflammatory target SFTPD was significantly decreased (**Figure 3**).

**Figure 3.**
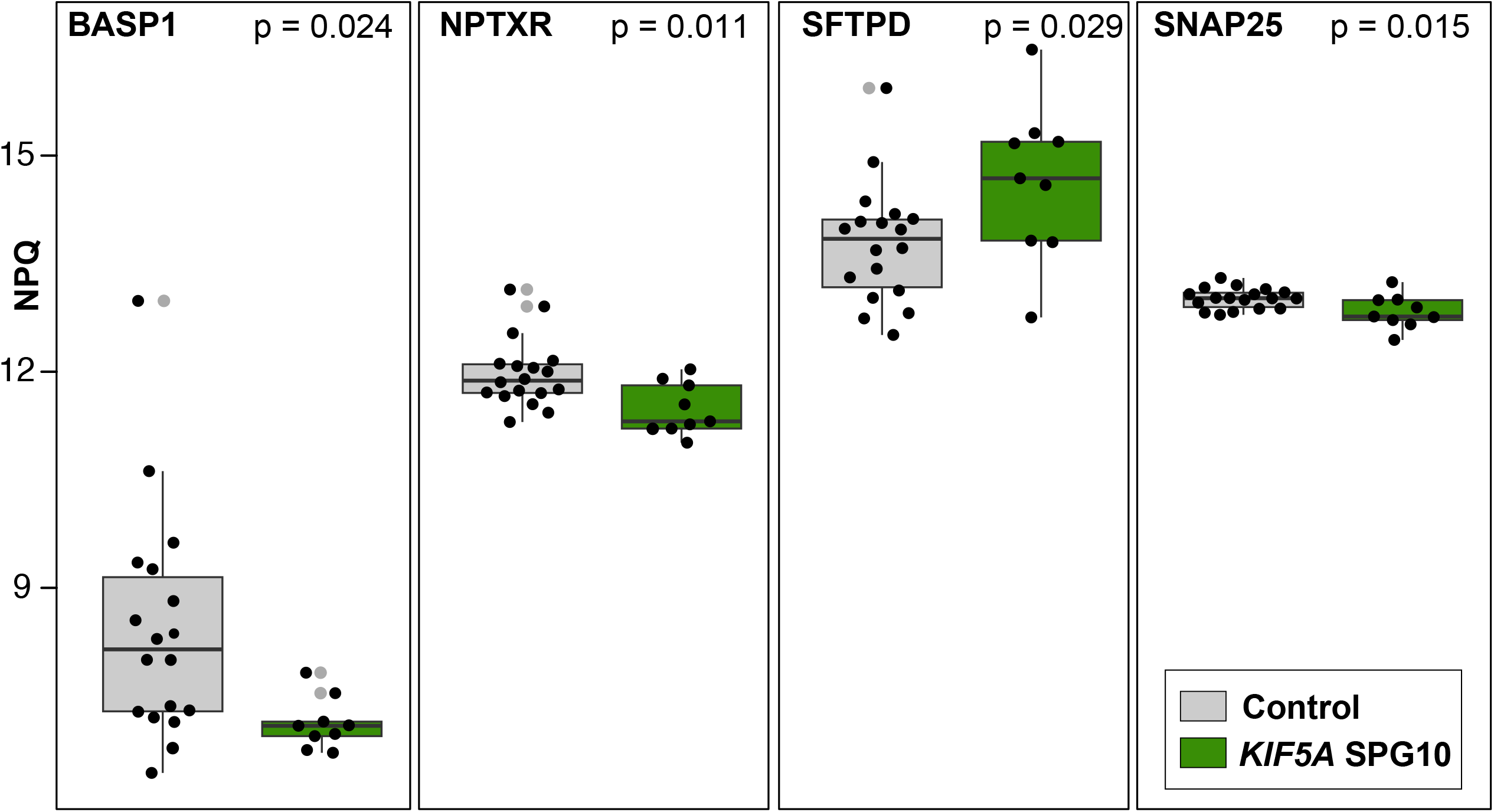
Comparison of NULISAseq biomarkers between control and *KIF5A*-linked SPG10. Box plots illustrating the distributions of targets in plasma of controls (n=18) and *KIF5A* SPG (n=9) participants with significant or marginally significant differences for proteins (BASP1, NPTXR, SNAP25 and SFTPD). The y-axis represents log_2_-tranformation of normalized read counts (NULISA Protein Quantification, NPQ). On each box, the central mark indicates the median, and the bottom and top edges of the box indicate the 25th and 75th percentiles, respectively. The whiskers extend to the most extreme data points not considered outliers, and the outliers are plotted individually using the grey dot symbol. The *p* values were determined using linear model. Significance determination was based on *p*-value < 0.05.

In the last step, we investigated the differentially expressed proteins in the ALS and SPG10 groups which could distinguish and possibly mirror the pathogenic mechanisms underlying axonopathy. We found significant changes in the BASP1, CNTN2 and NEFL proteins (**Figure 4**). In summary, the results highlighted significant proteins changes within the *KIF5A* ALS and SPG10 clinical spectrum. By identifying proteins that were differentially expressed under these conditions, the study offers valuable insights into physiological and pathological changes associated with *KIF5A* linked axonopathies. The differential expression of proteins identified in presymptomatic vs ALS suggests a dynamic relationship between neurodegeneration and early-stage alterations (**Figure S1**), providing new avenues to gain understanding how axon damage develops.

**Figure 4.**
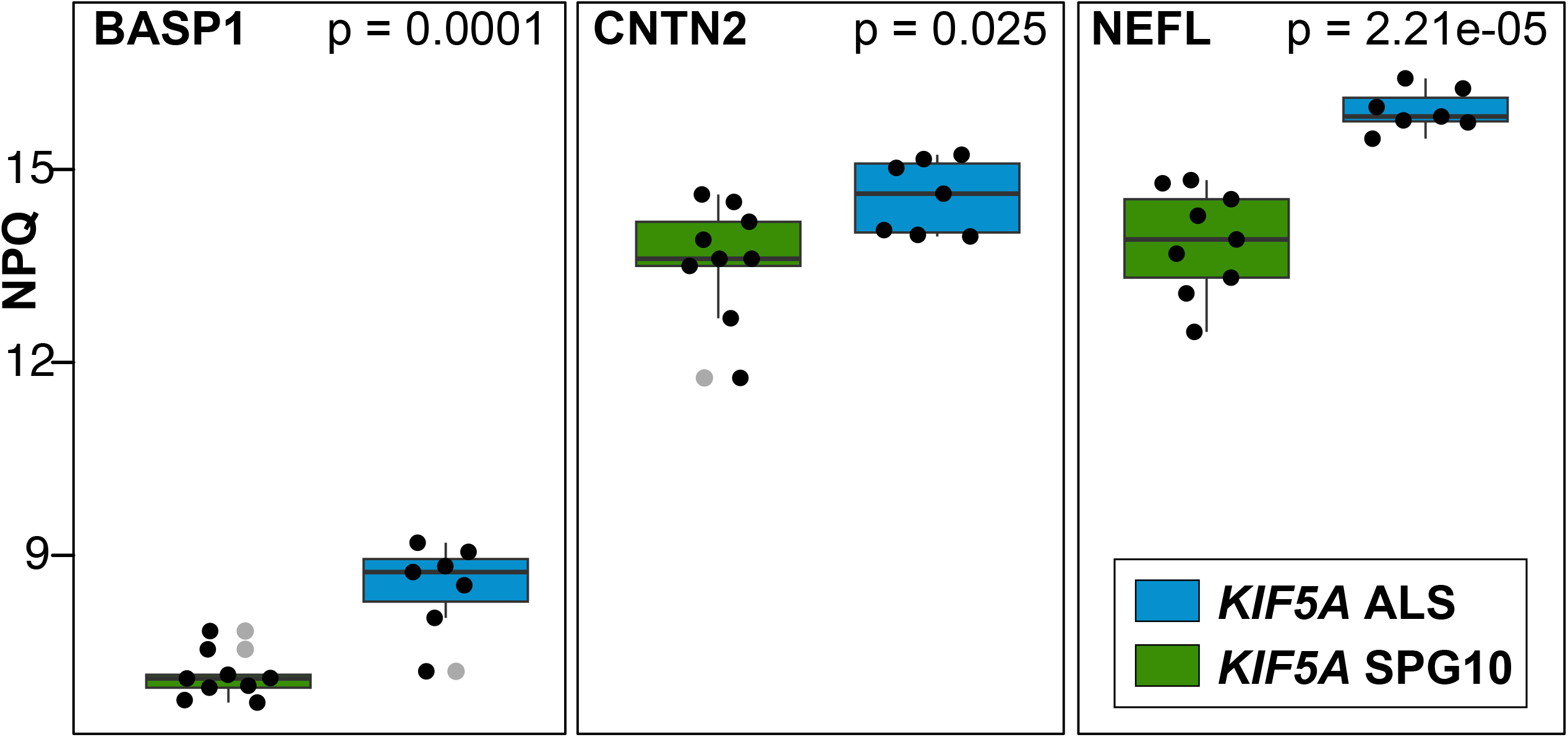
Comparison of NULISAseq biomarkers between *KIF5A* ALS and SPG10. Box plots illustrating the distributions of targets in plasma of KIF5A ALS (n=7) and KIF5A SPG (n=9) participants with significant or marginally significant differences for proteins (BASP1, CNTN2 and NEFL). The y-axis represents log_2_-tranformation of normalized read counts (NULISA Protein Quantification, NPQ). On each box, the central mark indicates the median, and the bottom and top edges of the box indicate the 25th and 75th percentiles, respectively. The whiskers extend to the most extreme data points not considered outliers, and the outliers are plotted individually using the grey dot symbol. The *p* values were determined using linear model. Significance determination was based on *p*-value < 0.05.

## Discussion

### Multiplex NULISA™ platform offers a promising approach for the study of neurodegenerative disorders

So far, blood-based biomarkers were performed using non-targeted liquid chromatography-tandem mass spectrometry techniques (LC-MS/MS). However, they are limited by the broad dynamic ranges of protein concentrations in plasma or serum where the detection of subtle variations is impeded by more abundant proteins, such as albumins and immunoglobulins.^27, 28^ Here we adopted a pioneered technology, a far more sensitive multiplex NULISA™ platform and focused on a defined set of central nervous system proteins in plasma samples that can be detected with attomolar sensitivity.^22^ Notably, neurodegenerative diseases (ALS, SPG, AD, PD) that affect axons often exhibit common phenotypic features, such as impairments in axonal transport or a “dying-back” process that begins at the distal synaptic terminals. In contrast, mutations in one gene can lead to different neurodegenerative disorders. To better understand these phenomena, it is crucial to examine axonopathies from both disease-specific and broader cell type perspectives. The type 1 kinesin (KIF5A) is a prime candidate for such an approach. KIF5A is a very prominent motor protein mediating anterograde microtubule-based transport in neurons, partly redundant with two KIF5 paralogues, various type 3 kinesin motors and potentially some other kinesin classes, such as type 2 kinesins.^29-31^

The loss of function of KIF5A in neurons would be expected to negatively impact on the provision with life-sustaining materials and organelles resulting in loss of axonal homeostasis and gradual degeneration; but it remains poorly explained as to why this affects primarily certain sub-types of neurons which then lead to disease-specific symptoms.^32, 33^ Similarly, we lack clear explanations for why different mutant alleles in the same motor protein, KIF5A, give rise to distinct neurodegenerative disease (**Figure 1A**). Some hypotheses arise from molecular analyses of KIF5A C-terminal ALS-linked high penetrant variants suggest these variants cause a gain of function rather than a loss. However, this does not clarify the specific pathological outcomes associated with these variants.^17, 19, 34-36^ We believe that new insights may emerge from blood plasma proteomics studies aimed at identifying possible molecular features that differ between SPG10- and ALS-linked *KIF5A* variants.

A few key measures were implemented to further refine our study: Firstly, the study was largely driven by international collaboration, which enabled the collection of plasma and critical clinical data from an exceptionally diverse cohort of recruited patients. Secondly, we ensured that *KIF5A* mutation carriers were compared to with age- and sex-matched healthy controls. Thirdly, the use of DNA barcodes on the NULISA™ platform facilitated multiplex proteomics, enabling us to run plasma samples from both patient and control groups together to minimize experimental variation in protein level differences. The data we obtained with the NULISA™ platform clearly revealed robust differences in protein expression between the compared groups, suggesting the platform’s future potential as diagnostic tool for KIF5A-linked SPG10, ALS patients and beyond. Moreover, we firmly believe that these findings could offer valuable mechanistic insights into neurodegenerative diseases and their progression. A critical question in this context is why proteins appear in the blood at varying levels, as it may reflect cell type-specific degrees of deterioration and/or the strength of target protein expression. For instance, we observed high levels of NEFL in one patient group and not the other, although axons affected in both groups are partly overlapping and all of them are enriched with neurofilaments.^37^ We believe that striving to understand such unfathomable observations like these opens up important new avenues of enquiry for the investigation of these diseases.

### ALS-linked KIF5A mutations correlate with a distinct plasma proteome

Our use of targeted multiplex plasma proteomics to identify brain-derived proteins revealed consistent protein signatures across multiple independent samples as clearly illustrated by the comparisons of plasma from *KIF5A* ALS patients to age-matched healthy controls (**Figure 2**). Strongest differences were observed for a number of genes with a diverse range of functions: upregulation of the APP fragment Aβ38 has previously been correlated with slowed cognitive decline in AD;^38, 39^ and the γ-secretase PSEN1 is linked to AD (OMIM #607822), FTD (#600274) and Pick disease (#172700); proteins with cytoskeletal links include the actin regulator CNTN2 (linked to epilepsy; #615400), the neurofilament subunit NEFL (linked to CMT; #617882, #607734, #607684) and the tau fragments pTau-181 and pTau-231 (predictors of AD and cognitive decline;^40^ four proteins are known inflammation factors including the chemokine CXCL1, the cytokine IL15, vascular inflammatory SFTPD and the fatty acid transporter FABP3. Further proteins comprise the synaptic transmembrane protein NPTX1 (linked to ataxia; #620158); and the ATP-dependent DNA regulator RUVBL2.^41-45^ Notably, the presence of various factors linked to other neural disorders in the blood of ALS patients may reflect the heterogeneity in clinical symptoms and disease progression.

The NEFL results are in line with previous report showing no or modest increase in SPG10 patients^46^, whereas phosphorylated neurofilament heavy chain was prominently elevated in serum and CSF of sporadic ALS patients compared with different SPG cases in a previous study.^47^ This further confirms the reproductivity of the NULISA assay. One possible explanation for the differences in NEFL could be the involvement of both upper and lower motor neurons, resulting in more widespread axonal damage with the release of larger amounts of NEFL, as occurs in typical ALS.^48, 49^

### SPG10-linked KIF5A mutations show a different signature from ALS patients

The plasma proteome of the SPG10 cohort identified four proteins that were significantly elevated compared to controls (**Figure 3**). Of these, only SFTPD was shared with the ALS cohort whereas the other proteins clearly deviated. These are all proteins localized to axonal terminals or synapses; they comprise the nerve growth and plasticity regulator BASP1^50^, the synaptic transmembrane protein NPTXR which is related to NPTX1 and considered a potential biomarker of synaptic dysfunction,^51^ and the t-SNARE SNAP25 pivotal for synaptic release and linked to neuromuscular myasthenic syndrome (#616330)^52, 53^. The upregulation of these factors might reflect the slow progression, resilience, and regenerative capacity in SPG10 patients. For example, the presence of BASP1 might be due to increases sprouting of regenerative collaterals ^50 54-57^, and enrichment of synaptic proteins might indicate synaptic loss or dysfunction involved in slow disease progression.

### Major conclusions and future perspectives

Our study is the first and largest comparative analysis of the *KIF5A* spectrum with patients from multiple continents and reveals promising correlations between *KIF5A* variants and underlying disease manifestations. To confirm results obtained here, longitudinal studies tracking proteomic changes over time would be a valuable next step, although such studies will face practical challenges given the slow disease progression reported in *KIF5A* patients, which can span 40 to 60 years. Nonetheless, the importance of such studies is underscored by our finding that presymptomatic individuals with known ALS-linked mutations exhibited down-regulation of several proteins (Aβ38, Aβ40, CCL11, GFAP, NEFL, SMOC1, VCAM1, VGF) compared to the symptomatic phase of the disease where elevated levels of Aβ38 and NEFL were prominent, likely indicators of axon degeneration (**Figure S1**). Therefore, to enable reliable early diagnoses, we need better descriptions of such dynamic changes over time. It is also important to note that identifying KIF5A patients for our study was a major effort, requiring outreach to numerous neuromuscular units worldwide. An international registry for patients with variants in the *KIF5A* gene would significantly facilitate more systematic studies, which could be complemented by repeated neuroimaging to establish closer correlations between observed morphological pathologies and blood proteins. We fully recognize the challenges of cohort size and validation, particularly given the rarity of *KIF5A*-related disorders. However, rather than a limitation, this can be viewed as an opportunity—our dataset provides the first insights into plasma proteomic changes in this population, establishing a foundation for future basic cell biology and translational studies. While an independent validation cohort was not feasible, our work aligns with a well-recognized precedent in rare neurodegenerative research, where numerous published studies, including those utilizing N=1 design, have made significant contributions. Such underpowered studies are increasingly valued in the field, particularly in motor neuron diseases (ALS, SPG10, CMT), as they support the growing emphasis on personalized medicine. Furthermore, the diagnostic accuracy of the applied method was assessed through a bootstrap procedure. The AUC calculations were performed for proteins with p < 0.05 across all comparisons. The resulting analysis identified targets that consistently showed significance, providing evidence that these targets are robust and not likely due to chance. In the context of rare diseases or complex comorbid conditions like *KIF5A* spectrum disorders, large-scale trials are often impractical or even inappropriate, highlighting the importance of smaller, well-characterized datasets in driving research and clinical progress. Our findings could further play a critical role in shaping natural history studies and guiding the design of clinical trials and therapies for *KIF5A*-linked multifactorial disorders.

## Data availability

Data used in this study are available within the article. Additional data supporting the information may be available upon reasonable request.

## Acknowledgements

We thank patients for their donation of biofluids. We would like to thank members of Jiang, Faundez laboratories at Emory, as well as the colleagues at Emory ALS center and the Emory proteomics core for helpful discussions. We would like to extend our gratitude to the Alamar Technology Access Program team for their valuable inputs.

## Supplementary Material

**Table S1. Comparison of target performance across groups using AUC metrics from bootstrap analysis**.

**Figure S1.**
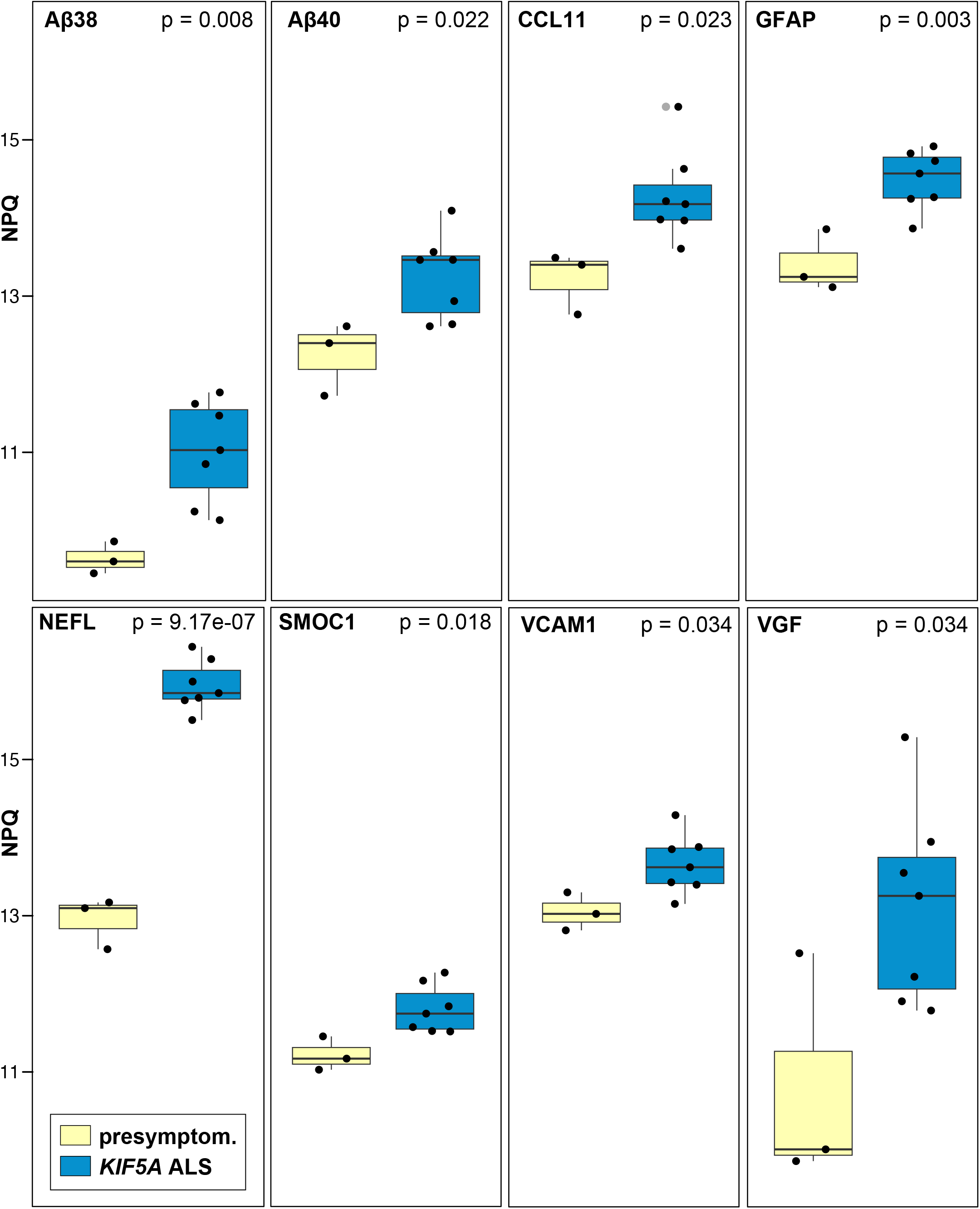
Comparison of NULISAseq biomarkers between presymptomatic and *KIF5A* ALS. Box plots illustrating the distributions of targets in plasma of KIF5A ALS (n=7) and presymptom (presymptomatic, n=3) participants with significant or marginally significant differences for proteins (Aβ38, Aβ40, CCL11, VCAM1, GFAP, NEFL, SMOC1, and VGF). The y-axis represents log_2_-tranformation of normalized read counts (NULISA Protein Quantification, NPQ). On each box, the central mark indicates the median, and the bottom and top edges of the box indicate the 25th and 75th percentiles, respectively. The whiskers extend to the most extreme data points not considered outliers, and the outliers are plotted individually using the grey dot symbol. The *p* values were determined using linear model. Significance determination was based on *p*-value < 0.05.

**Figure S2.**
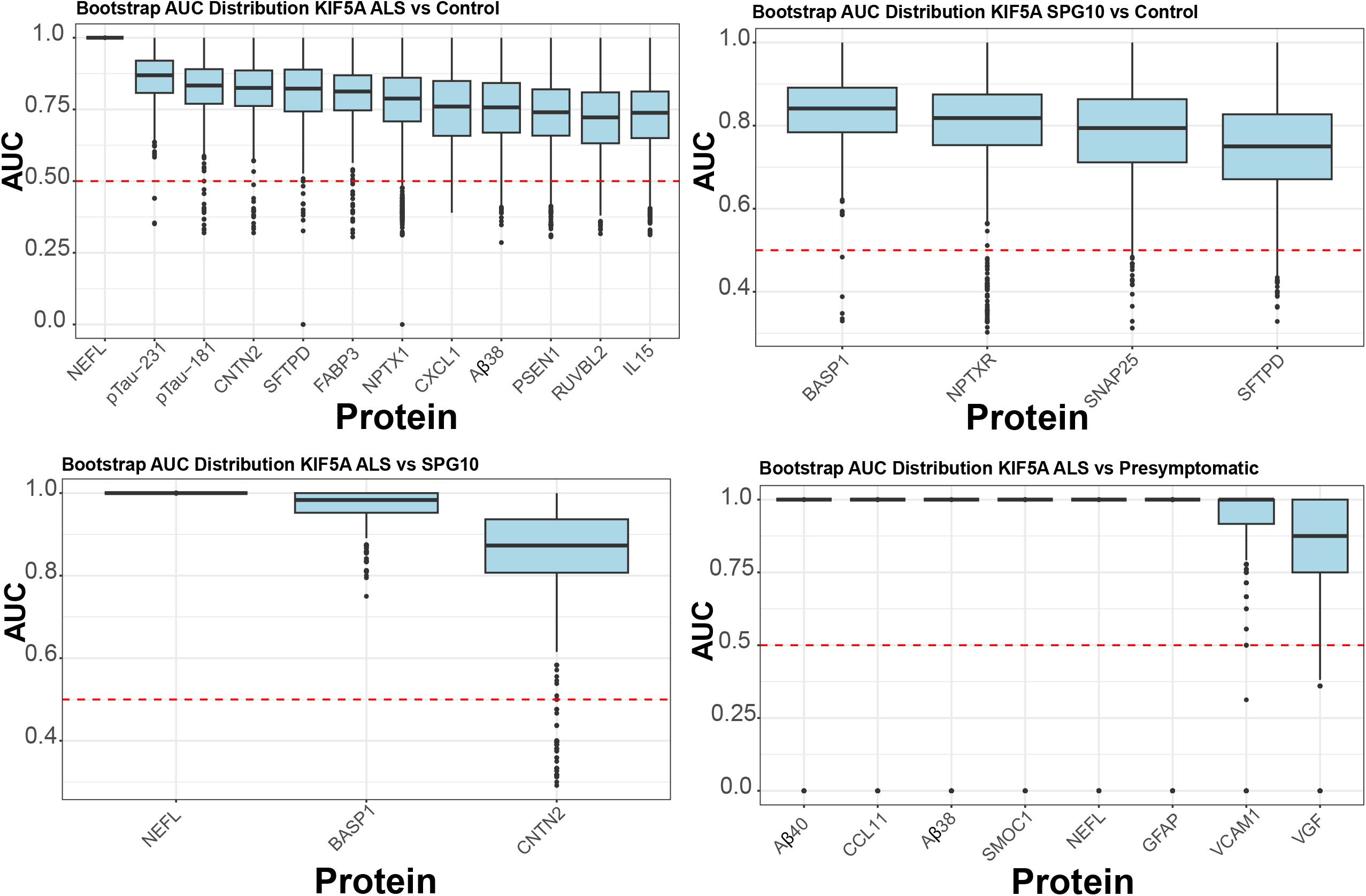
Box plot comparing accuracy assessment statistic area under the receiver operating characteristic curve (AUC) for different *KIF5A* groups. Box plots of the AUC values showing median, inter-quartile ranges and total ranges from the 1000 bootstrap iterations. The red dashed line indicates the mean AUC derived from random distribution.

## References

1. Hirokawa N. Kinesin and dynein superfamily proteins and the mechanism of organelle transport. Science. Jan 23 1998;279(5350):519–26. doi:10.1126/science.279.5350.519

2. Brenner D, Yilmaz R, Muller K, et al. Hot-spot KIF5A mutations cause familial ALS. Brain. Mar 1 2018;141(3):688–697. doi:10.1093/brain/awx370

3. Crimella C, Baschirotto C, Arnoldi A, et al. Mutations in the motor and stalk domains of KIF5A in spastic paraplegia type 10 and in axonal Charcot-Marie-Tooth type 2. Clin Genet. Aug 2012;82(2):157–64. doi:10.1111/j.1399-0004.2011.01717.x

4. Duis J, Dean S, Applegate C, et al. KIF5A mutations cause an infantile onset phenotype including severe myoclonus with evidence of mitochondrial dysfunction. Ann Neurol. Oct 2016;80(4):633–7. doi:10.1002/ana.24744

5. Nicolas A, Kenna KP, Renton AE, et al. Genome-wide Analyses Identify KIF5A as a Novel ALS Gene. Neuron. Mar 21 2018;97(6):1267–1288. doi:10.1016/j.neuron.2018.02.027

6. Reid E, Kloos M, Ashley-Koch A, et al. A kinesin heavy chain (KIF5A) mutation in hereditary spastic paraplegia (SPG10). Am J Hum Genet. Nov 2002;71(5):1189–94. doi:10.1086/344210

7. Rydzanicz M, Jagla M, Kosinska J, et al. KIF5A de novo mutation associated with myoclonic seizures and neonatal onset progressive leukoencephalopathy. Clin Genet. May 2017;91(5):769–773. doi:10.1111/cge.12831

8. Reid E, Dearlove AM, Rhodes M, Rubinsztein DC. A new locus for autosomal dominant “pure” hereditary spastic paraplegia mapping to chromosome 12q13, and evidence for further genetic heterogeneity. Am J Hum Genet. Sep 1999;65(3):757–63. doi:10.1086/302555

9. Goizet C, Boukhris A, Mundwiller E, et al. Complicated forms of autosomal dominant hereditary spastic paraplegia are frequent in SPG10. Hum Mutat. Feb 2009;30(2):E376–85. doi:10.1002/humu.20920

10. Kaji S, Kawarai T, Miyamoto R, et al. Late-onset spastic paraplegia type 10 (SPG10) family presenting with bulbar symptoms and fasciculations mimicking amyotrophic lateral sclerosis. J Neurol Sci. May 15 2016;364:45–9. doi:10.1016/j.jns.2016.03.001

11. de Fuenmayor-Fernandez de la Hoz CP, Hernandez-Lain A, Olive M, Sanchez-Calvin MT, Gonzalo-Martinez JF, Dominguez-Gonzalez C. Adult-onset distal spinal muscular atrophy: a new phenotype associated with KIF5A mutations. Brain. Dec 1 2019;142(12):e66. doi:10.1093/brain/awz317

12. Kenna KP, van Doormaal PT, Dekker AM, et al. NEK1 variants confer susceptibility to amyotrophic lateral sclerosis. Nat Genet. Sep 2016;48(9):1037–42. doi:10.1038/ng.3626

13. Saez-Atienzar S, Dalgard CL, Ding J, et al. Identification of a pathogenic intronic KIF5A mutation in an ALS-FTD kindred. Neurology. Dec 1 2020;95(22):1015–1018. doi:10.1212/WNL.0000000000011064

14. Yoo KS, Lee K, Oh JY, et al. Postsynaptic density protein 95 (PSD-95) is transported by KIF5 to dendritic regions. Mol Brain. Nov 21 2019;12(1):97. doi:10.1186/s13041-019-0520-x

15. Wang L, Brown A. A hereditary spastic paraplegia mutation in kinesin-1A/KIF5A disrupts neurofilament transport. Mol Neurodegener. Nov 18 2010;5:52. doi:10.1186/1750-1326-5-52

16. Fukuda Y, Pazyra-Murphy MF, Silagi ES, et al. Binding and transport of SFPQ-RNA granules by KIF5A/KLC1 motors promotes axon survival. J Cell Biol. Jan 4 2021;220(1) doi:10.1083/jcb.202005051

17. Baron DM, Fenton AR, Saez-Atienzar S, et al. ALS-associated KIF5A mutations abolish autoinhibition resulting in a toxic gain of function. Cell Rep. Apr 5 2022;39(1):110598. doi:10.1016/j.celrep.2022.110598

18. Ebbing B, Mann K, Starosta A, et al. Effect of spastic paraplegia mutations in KIF5A kinesin on transport activity. Hum Mol Genet. May 1 2008;17(9):1245–52. doi:10.1093/hmg/ddn014

19. Nakano J, Chiba K, Niwa S. An ALS-associated KIF5A mutant forms oligomers and aggregates and induces neuronal toxicity. Genes Cells. Jun 2022;27(6):421–435. doi:10.1111/gtc.12936

20. Pant DC, Parameswaran J, Rao L, et al. ALS-linked KIF5A DeltaExon27 mutant causes neuronal toxicity through gain-of-function. EMBO Rep. Aug 3 2022;23(8):e54234. doi:10.15252/embr.202154234

21. Brooks BR. El Escorial World Federation of Neurology criteria for the diagnosis of amyotrophic lateral sclerosis. Subcommittee on Motor Neuron Diseases/Amyotrophic Lateral Sclerosis of the World Federation of Neurology Research Group on Neuromuscular Diseases and the El Escorial “Clinical limits of amyotrophic lateral sclerosis” workshop contributors. J Neurol Sci. Jul 1994;124 Suppl:96–107. doi:10.1016/0022-510x(94)90191-0

22. Feng W, Beer JC, Hao Q, et al. NULISA: a proteomic liquid biopsy platform with attomolar sensitivity and high multiplexing. Nat Commun. Nov 9 2023;14(1):7238. doi:10.1038/s41467-023-42834-x

23. Irwin KE, Sheth U, Wong PC, Gendron TF. Fluid biomarkers for amyotrophic lateral sclerosis: a review. Mol Neurodegener. Jan 24 2024;19(1):9. doi:10.1186/s13024-023-00685-6

24. Benatar M, Wuu J, Turner MR. Neurofilament light chain in drug development for amyotrophic lateral sclerosis: a critical appraisal. Brain. Jul 3 2023;146(7):2711–2716. doi:10.1093/brain/awac394

25. Benatar M, Wuu J, Andersen PM, Lombardi V, Malaspina A. Neurofilament light: A candidate biomarker of presymptomatic amyotrophic lateral sclerosis and phenoconversion. Ann Neurol. Jul 2018;84(1):130–139. doi:10.1002/ana.25276

26. Dulski J, Strongosky AJ, Al-Shaikh RH, Wszolek ZK. Expanding the spectrum of KIF5A mutations-case report of a large kindred with familial ALS and overlapping syndrome. Amyotroph Lateral Scler Frontotemporal Degener. May 2023;24(3-4):347–350. doi:10.1080/21678421.2022.2164204

27. Anderson NL, Polanski M, Pieper R, et al. The human plasma proteome: a nonredundant list developed by combination of four separate sources. Mol Cell Proteomics. Apr 2004;3(4):311–26. doi:10.1074/mcp.M300127-MCP200

28. Rzepinski L, Koslinski P, Kowalewski M, Koba M, Maciejek Z. Serum amino acid profiling in differentiating clinical outcomes of multiple sclerosis. Neurol Neurochir Pol. 2023;57(5):414–422. doi:10.5603/PJNNS.a2023.0054

29. Maday S, Twelvetrees AE, Moughamian AJ, Holzbaur EL. Axonal transport: cargo-specific mechanisms of motility and regulation. Neuron. Oct 22 2014;84(2):292–309. doi:10.1016/j.neuron.2014.10.019

30. Hirokawa N, Niwa S, Tanaka Y. Molecular motors in neurons: transport mechanisms and roles in brain function, development, and disease. Neuron. Nov 18 2010;68(4):610–38. doi:10.1016/j.neuron.2010.09.039

31. Kanai Y, Okada Y, Tanaka Y, Harada A, Terada S, Hirokawa N. KIF5C, a novel neuronal kinesin enriched in motor neurons. J Neurosci. Sep 1 2000;20(17):6374–84. doi:10.1523/JNEUROSCI.20-17-06374.2000

32. Smith G, Sweeney ST, O’Kane CJ, Prokop A. How neurons maintain their axons long-term: an integrated view of axon biology and pathology. Front Neurosci. 2023;17:1236815. doi:10.3389/fnins.2023.1236815

33. Prokop A. A common theme for axonopathies? The dependency cycle of local axon homeostasis. Cytoskeleton (Hoboken). Feb 2021;78(2):52–63. doi:10.1002/cm.21657

34. Carrington G, Fatima U, Caramujo I, Lewis T, Casas-Mao D, Peckham M. A multiscale approach reveals the molecular architecture of the autoinhibited kinesin KIF5A. J Biol Chem. Mar 2024;300(3):105713. doi:10.1016/j.jbc.2024.105713

35. Cozzi M, Magri S, Tedesco B, et al. Altered molecular and cellular mechanisms in KIF5A-associated neurodegenerative or neurodevelopmental disorders. Cell Death Dis. Sep 27 2024;15(9):692. doi:10.1038/s41419-024-07096-5

36. Pant DC, Verma S. Identifying novel response markers for spinal muscular atrophy revealed by targeted proteomics following gene therapy. Gene Ther. Jan 10 2025; doi:10.1038/s41434-025-00513-0

37. Prokop A. Cytoskeletal organization of axons in vertebrates and invertebrates. J Cell Biol. Jul 6 2020;219(7) doi:10.1083/jcb.201912081

38. Cullen N, Janelidze S, Palmqvist S, et al. Association of CSF Abeta(38) Levels With Risk of Alzheimer Disease-Related Decline. Neurology. Mar 1 2022;98(9):e958–e967. doi:10.1212/WNL.0000000000013228

39. Lewczuk P, Lukaszewicz-Zajac M, Kornhuber J, Mroczko B. Clinical significance of plasma candidate biomarkers of Alzheimer’s Disease. Neurol Neurochir Pol. 2024;58(4):363–379. doi:10.5603/pjnns.100675

40. Tissot C, Therriault J, Kunach P, et al. Comparing tau status determined via plasma pTau181, pTau231 and [(18)F]MK6240 tau-PET. EBioMedicine. Feb 2022;76:103837. doi:10.1016/j.ebiom.2022.103837

41. Colmorten KB, Nexoe AB, Sorensen GL. The Dual Role of Surfactant Protein-D in Vascular Inflammation and Development of Cardiovascular Disease. Front Immunol. 2019;10:2264. doi:10.3389/fimmu.2019.02264

42. Gomez de San Jose N, Massa F, Halbgebauer S, Oeckl P, Steinacker P, Otto M. Neuronal pentraxins as biomarkers of synaptic activity: from physiological functions to pathological changes in neurodegeneration. J Neural Transm (Vienna). Feb 2022;129(2):207–230. doi:10.1007/s00702-021-02411-2

43. Korbecki J, Gassowska-Dobrowolska M, Wojcik J, et al. The Importance of CXCL1 in Physiology and Noncancerous Diseases of Bone, Bone Marrow, Muscle and the Nervous System. Int J Mol Sci. Apr 11 2022;23(8) doi:10.3390/ijms23084205

44. Nguyen HC, Bu S, Nikfarjam S, et al. Loss of fatty acid binding protein 3 ameliorates lipopolysaccharide-induced inflammation and endothelial dysfunction. J Biol Chem. Mar 2023;299(3):102921. doi:10.1016/j.jbc.2023.102921

45. Perera PY, Lichy JH, Waldmann TA, Perera LP. The role of interleukin-15 in inflammation and immune responses to infection: implications for its therapeutic use. Microbes Infect. Mar 2012;14(3):247–61. doi:10.1016/j.micinf.2011.10.006

46. Andreasson M, Lagerstedt-Robinson K, Samuelsson K, et al. Altered CSF levels of monoamines in hereditary spastic paraparesis 10: A case series. Neurol Genet. Aug 2019;5(4):e344. doi:10.1212/NXG.0000000000000344

47. Simonini C, Zucchi E, Bedin R, et al. CSF Heavy Neurofilament May Discriminate and Predict Motor Neuron Diseases with Upper Motor Neuron Involvement. Biomedicines. Nov 5 2021;9(11) doi:10.3390/biomedicines9111623

48. Mills KR. Characteristics of fasciculations in amyotrophic lateral sclerosis and the benign fasciculation syndrome. Brain. Nov 2010;133(11):3458–69. doi:10.1093/brain/awq290

49. Brugman F, Veldink JH, Franssen H, et al. Differentiation of hereditary spastic paraparesis from primary lateral sclerosis in sporadic adult-onset upper motor neuron syndromes. Arch Neurol. Apr 2009;66(4):509–14. doi:10.1001/archneurol.2009.19

50. Chung D, Shum A, Caraveo G. GAP-43 and BASP1 in Axon Regeneration: Implications for the Treatment of Neurodegenerative Diseases. Front Cell Dev Biol. 2020;8:567537. doi:10.3389/fcell.2020.567537

51. Dulewicz M, Kulczynska-Przybik A, Borawska R, Slowik A, Mroczko B. The neuronal pentraxin receptor (NPTXR) as a candidate biomarker of synaptic dysfunction in mild cognitive impairment. Alzheimer’s & Dementia. 2023;19(S2):e063018. doi:10.1002/alz.063018

52. Zhou J, Wade SD, Graykowski D, et al. The neuronal pentraxin Nptx2 regulates complement activity and restrains microglia-mediated synapse loss in neurodegeneration. Sci Transl Med. Mar 29 2023;15(689):eadf0141. doi:10.1126/scitranslmed.adf0141

53. Batista AFR, Martinez JC, Hengst U. Intra-axonal Synthesis of SNAP25 Is Required for the Formation of Presynaptic Terminals. Cell Rep. Sep 26 2017;20(13):3085–3098. doi:10.1016/j.celrep.2017.08.097

54. Filosto M, Piccinelli SC, Palmieri I, et al. A Novel Mutation in the Stalk Domain of KIF5A Causes a Slowly Progressive Atypical Motor Syndrome. J Clin Med. Dec 22 2018;8(1) doi:10.3390/jcm8010017

55. Schule R, Kremer BP, Kassubek J, et al. SPG10 is a rare cause of spastic paraplegia in European families. J Neurol Neurosurg Psychiatry. May 2008;79(5):584–7. doi:10.1136/jnnp.2007.137596

56. Meyyazhagan A, Orlacchio A. Hereditary Spastic Paraplegia: An Update. Int J Mol Sci. Feb 1 2022;23(3) doi:10.3390/ijms23031697

57. Martinello C, Panza E, Orlacchio A. Hereditary spastic paraplegias proteome: common pathways and pathogenetic mechanisms. Expert Rev Proteomics. Jul-Dec 2023;20(7-9):171–188. doi:10.1080/14789450.2023.2260952

58. de Boer EMJ, van Rheenen W, Goedee HS, et al. Genotype-phenotype correlations of KIF5A stalk domain variants. Amyotroph Lateral Scler Frontotemporal Degener. Nov 2021;22(7-8):561–570. doi:10.1080/21678421.2021.1907412

59. He J, Liu X, Tang L, Zhao C, He J, Fan D. Whole-exome sequencing identified novel KIF5A mutations in Chinese patients with amyotrophic lateral sclerosis and Charcot-Marie-Tooth type 2. J Neurol Neurosurg Psychiatry. Mar 2020;91(3):326–328. doi:10.1136/jnnp-2019-320483

60. Naruse H, Ishiura H, Mitsui J, et al. Splice-site mutations in KIF5A in the Japanese case series of amyotrophic lateral sclerosis. Neurogenetics. Mar 2021;22(1):11–17. doi:10.1007/s10048-020-00626-1

