## Supplementary material for "Targeted plasma proteomics uncover novel proteins associated with *KIF5A*-linked SPG10 and ALS spectrum disorders": Table S1

| Comparison | Protein | MeanAUC | MedianAUC | CI_Lower | CI_Upper | P_Value | Significant |
|---|---|---|---|---|---|---|---|
| ALS_vs_Asymp | Aβ38 | 0.981 | 1 | 1 | 1 | 0.019 | TRUE |
| ALS_vs_Asymp | SMOC1 | 0.979 | 1 | 1 | 1 | 0.021 | TRUE |
| ALS_vs_Asymp | VCAM1 | 0.93033921 | 1 | 0.38080357 | 1 | 0.026 | TRUE |
| ALS_vs_Asymp | CCL11 | 0.972 | 1 | 0 | 1 | 0.028 | TRUE |
| ALS_vs_Asymp | Aβ40 | 0.97 | 1 | 0 | 1 | 0.03 | TRUE |
| ALS_vs_Asymp | NEFL | 0.968 | 1 | 0 | 1 | 0.032 | TRUE |
| ALS_vs_Asymp | GFAP | 0.966 | 1 | 0 | 1 | 0.034 | TRUE |
| ALS_vs_Asymp | VGF | 0.84490206 | 0.9047619 | 0 | 1 | 0.068 | FALSE |
| ALS_vs_CTL | NEFL | 1 | 1 | 1 | 1 | 0 | TRUE |
| ALS_vs_CTL | pTau-231 | 0.85826744 | 0.86764706 | 0.68411915 | 0.97794118 | 0.004 | TRUE |
| ALS_vs_CTL | CNTN2 | 0.81565472 | 0.82 | 0.63482143 | 0.96034799 | 0.009 | TRUE |
| ALS_vs_CTL | pTau-181 | 0.82729957 | 0.84 | 0.624875 | 0.98814286 | 0.015 | TRUE |
| ALS_vs_CTL | SFTPD | 0.80799431 | 0.81818182 | 0.56975 | 0.98414502 | 0.019 | TRUE |
| ALS_vs_CTL | FABP3 | 0.7922907 | 0.80555556 | 0.49980159 | 0.958375 | 0.029 | TRUE |
| ALS_vs_CTL | CXCL1 | 0.74819859 | 0.75 | 0.46316235 | 1 | 0.04 | TRUE |
| ALS_vs_CTL | Aβ38 | 0.74345113 | 0.75 | 0.42837302 | 0.97619048 | 0.062 | FALSE |
| ALS_vs_CTL | NPTX1 | 0.76777745 | 0.78571429 | 0.38454861 | 0.9800614 | 0.066 | FALSE |
| ALS_vs_CTL | RUVBL2 | 0.71212034 | 0.72222222 | 0.39992647 | 0.96000794 | 0.081 | FALSE |
| ALS_vs_CTL | PSEN1 | 0.72060395 | 0.74 | 0.40475315 | 0.95614035 | 0.088 | FALSE |
| ALS_vs_CTL | IL15 | 0.7169851 | 0.73 | 0.38666667 | 0.96039041 | 0.098 | FALSE |
| ALS_vs_SPG10 | NEFL | 1 | 1 | 1 | 1 | 0 | TRUE |
| ALS_vs_SPG10 | BASP1 | 0.96770135 | 0.98412698 | 0.84114719 | 1 | 0.001 | TRUE |
| ALS_vs_SPG10 | CNTN2 | 0.8434514 | 0.86923077 | 0.390625 | 1 | 0.046 | TRUE |
| SPG10_vs_CTL | BASP1 | 0.83105812 | 0.83571429 | 0.65783626 | 0.9691358 | 0.003 | TRUE |
| SPG10_vs_CTL | SNAP25 | 0.78685174 | 0.8 | 0.5272488 | 0.96919312 | 0.015 | TRUE |
| SPG10_vs_CTL | NPTXR | 0.79975381 | 0.81481481 | 0.41303485 | 0.98571429 | 0.044 | TRUE |
| SPG10_vs_CTL | SFTPD | 0.739339 | 0.75 | 0.43173701 | 0.94744964 | 0.051 | FALSE |
